# Morphological facilitators for vocal learning explored through the syrinx anatomy of a basal hummingbird

**DOI:** 10.1101/2020.01.11.902734

**Authors:** Amanda Monte, Alexander F. Cerwenka, Bernhard Ruthensteiner, Manfred Gahr, Daniel N. Düring

## Abstract

Vocal learning is a rare evolutionary trait that evolved independently in three avian clades: songbirds, parrots, and hummingbirds. Although the anatomy and mechanisms of sound production in songbirds are well understood, little is known about the hummingbird’s vocal anatomy. We use high-resolution micro-computed tomography (μCT) and microdissection to reveal the three-dimensional structure of the syrinx, the vocal organ of the black jacobin (Florisuga fusca), a phylogenetically basal hummingbird species. We identify three unique features of the black jacobin’s syrinx: (i) a shift in the position of the syrinx to the outside of the thoracic cavity and the related loss of the sterno-tracheal muscle, (ii) complex intrinsic musculature, oriented dorso-ventrally, and (iii) ossicles embedded in the medial vibratory membranes. Their syrinx morphology allows vibratory decoupling, precise control of complex acoustic parameters, and a large redundant acoustic space that may be key biomechanical factors facilitating the occurrence of vocal production learning.

## Introduction

Vocal learning in birds -- the rare ability to acquire new sounds from the environment -- holds striking parallels with speech acquisition in humans (Marler, 1970). Vocal learning evolved independently in songbirds (suborder *Oscines*) (Bottjer et al., 1985; Nottebohm et al., 1976), parrots (order *Psittaciformes*) (Jarvis and Mello, 2000) and hummingbirds (family *Trochilidae*) (Baptista and Schuchmann, 1990) due to convergent neurological shifts (Jarvis, 2007). Thus, the brains of avian vocal learners are uniquely specialized, unlike non-vocal-learner species, to perceive, produce and memorize sounds (Gahr, 2000; Nottebohm et al., 1976; Paton et al., 1981). However, the pressures underlying this independent convergence remain unknown and attempts to explain the evolution of vocal learning face challenges from the divergences in the life histories of the vocal learners (Jarvis, 2006; Nowicki and Searcy, 2014).

Efforts to understand vocal learning have concentrated on the neural processes that modulate vocal output with little regard to the integral part given by the biomechanics of sound production in the vocal organ (Düring and Elemans, 2016). The vocal organ in birds is the syrinx (Clarke et al., 2016; King, 1989), an avian novelty optimized for birds’ particularly long air tracts (Riede et al., 2019). The syrinx is located where the trachea bifurcates into the bronchi and is suspended inside the interclavicular air sac (King, 1989). One or two pairs of vibrating tissues are present; depending on where these tissues are located, the syrinx can be classified as tracheal, tracheobronchial or bronchial (Smolker, 1947). The syrinx musculature is of two basic types: extrinsic musculature, which is attached outside of the syrinx at one end, and intrinsic musculature, which is attached to the syrinx at both ends (Ames, 1971; Gaunt and Gaunt, 1985). While every bird has extrinsic musculature, not all syrinxes have intrinsic musculature (Gaunt and Gaunt, 1985), which varies among birds (Ames, 1971; King, 1989). For example, gallinaceous species have none, songbird species have from three to five intrinsic muscle pairs (Ames, 1971; Gaunt, 1983), while parrot species mainly have two (Gaunt, 1983; Nottebohm, 1976).

Syrinx anatomy, in general, is highly variable among and consistent within higher-level taxa, to the extent that syrinxes have been used as a taxonomic tool for avian phylogenetic classification (Ames, 1971; Düring and Elemans, 2016; Suthers and Zollinger, 2008). Similarities in gross morphology and its implications for vocal production may help us to understand the morphological basis of vocal learning (Elemans et al., 2015; Gaunt, 1983). Thus, the presence of intrinsic musculature has been hypothesized as a prerequisite and not an adaptation for vocal learning (Gaunt, 1983; Mindlin and Laje, 2006), that is, all vocal learners should have intrinsic muscles, but not all species that have intrinsic muscles are vocal learners. Unlike extrinsic muscles, which move the syrinx as a unit (Gaunt, 1983; Mindlin and Laje, 2006), intrinsic muscles dissociate the control of tension from the control of amplitude, for example, which in turn affects pitch (Düring et al., 2017; Goller and Riede, 2013).

Recent studies indicate that musculature is just one of the variables that define the multi-dimensional parameter space that translates motor commands into vocal output (Amador and Mindlin, 2008; Düring et al., 2017; Düring and Elemans, 2016; Elemans et al., 2015). Many factors, such as syrinx’s morphology, physical interaction with the surrounding environment, and neuro-mechanic activity, contribute to the creation of a large acoustic space that is highly redundant (Elemans et al., 2015). This redundancy allows specific vocal parameters, such as pitch, to be achieved by multiple combinations of, for example, expiratory air pressure and muscle activity (Elemans et al., 2015). The availability of multiple motor commands for a certain acoustic target may simplify the trial-and-error learning process and is hypothesized as necessary for the development of vocal learning (Elemans et al., 2015).

To approach vocal learning from the biomechanical perspective, the syrinxes of vocal learners need to be systematically compared. Among avian vocal learners, hummingbirds are the most basal taxon and phylogenetic distant from parrots and songbirds (Baptista and Schuchmann, 1990; Gahr, 2000; Jarvis et al., 2000; Prum et al., 2015), and the only group in which not all species have the ability of vocal learning (Gahr, 2000; Williams and Houtman, 2008). The acoustic features of their vocalizations vary substantially within the group (Ferreira et al., 2006; Ficken et al., 2000), ranging from simple vocalizations to acoustic performances that are above the known perceptual limits of birds (Duque et al., 2018; Olson et al., 2018). Currently, we lack a detailed description of the hummingbird syrinx and, therefore, insights into the biomechanics of hummingbirds’ peculiar vocalizations.

Here we use micro-computed tomography (μCT) and microdissection to resolve the detailed structure of both skeletal elements and vibratory soft tissues of the black jacobin (*Florisuga fusca*) syrinx. The black jacobin belongs to the clade Topaze (subfamily *Topazini*), the most basal among hummingbirds (McGuire et al., 2014). It can vocalize on high F0 with harmonics over the human audible range (Olson et al., 2018). Our results provide fundamental insights into the biomechanics of sound production in hummingbirds and the anatomical factors driving or facilitating the emergence of vocal learning in birds.

## Results

### General anatomy of the black jacobin’s syrinx

The black jacobin has a tracheobronchial syrinx that is located where the trachea bifurcates into the two primary bronchi, approximately 9.4 mm far from the heart and outside of the thoracic cavity. The trachea is a long, funnel-shaped tube that extends along the cervical column whose diameter is reduced to around ¼ of its original size when proximal to the syrinx. From the syrinx on, the bronchi run parallel for about 1.3 mm, separating when inside the thorax. Parts of the trachea and bronchi, and the syrinx, are tightly packed by a multi-layered membrane, most likely an evagination of the clavicular air sac membrane (Zusi, 2013).

### The syringeal skeleton of the black jacobin

The syringeal skeleton of the black jacobin is composed of cartilaginous tracheal rings, a solid tympanum and modified bronchial half-rings, two of which are partially ossified (Fig. 1A and 1C). The trachea consists of complete cartilaginous rings (T1 to Tn), each of which is thinner in its dorsal part and wider towards the tympanum. The tympanum is a cylindrical bone likely formed by the fusion of tracheal and bronchial rings; this fusion forms the tympanum in other tracheobronchial syrinxes, for example, that of the zebra finch syrinx (Düring et al., 2013). Internally, the tympanum body is a relatively uniform tube with an ossified lamella in its caudal part that projects medially into the air passage. Externally, the ventral part of the tympanum body presents a squared expansion that, together with the internal lamella, constitutes the pessulus. The pessulus separates a symmetrical pair of horizontal ridges that delimit medially the two main craniocaudal concavities to which muscles are attached (Fig. 1B). The U-shaped dorsal part of the tympanum is formed by two solid expansions in each of the lateral caudal edges and a medial concavity that extends horizontally along the entire surface as a muscle attachment site. In the caudomedial part, a pair of rounded bones, the tympanic ossicles (*ossicula tympanica*), are embedded in an extension of the most medial part of the vibratory tissues (Fig. 1D).

**Fig 1.**
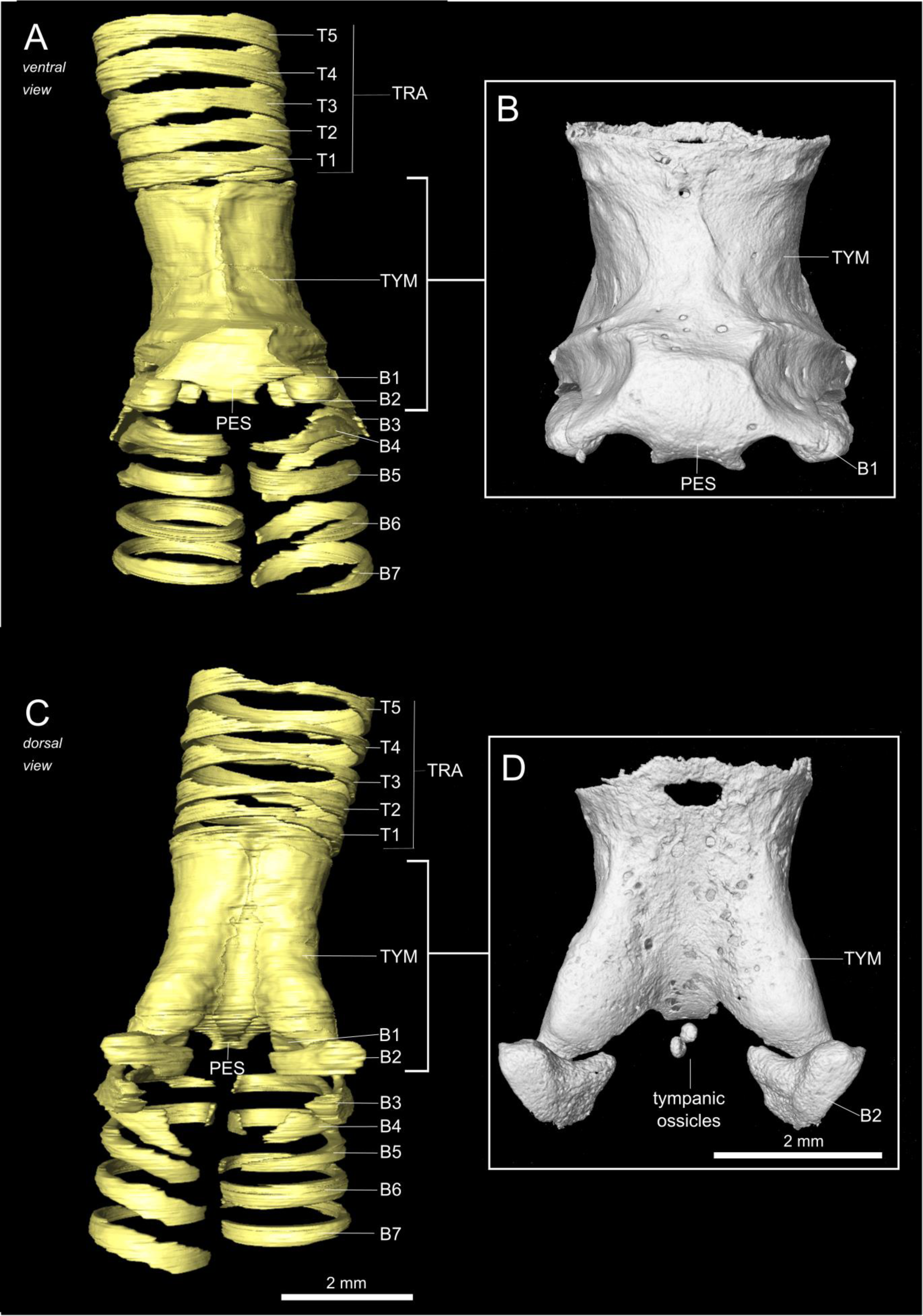
The syringeal skeleton of the black jacobin. 3D visualizations of µCT data. A and C, surface renderings of the contrasted sample; B and D, volume renderings of the non-contrasted sample. A and B, ventral view; C and D, dorsal view. In D, the tympanic ossicles can be seen. T1 to T5, tracheal rings; TYM, tympanum; B1 to B7, bronchial half-rings and PES, pessulus.

Caudally to the tympanum, bronchial half-rings (B1 to Bn) extend for around 12.3 mm, until reaching the lungs. Only the first two pairs are partially ossified (B1 and B2); the others are cartilaginous. The first pairs (B1 to B3) are highly modified compared to the other bronchial half-rings (Fig. 1A and 1B). Each ring of the B1 pair is composed of a ventral spherical ossification, and a cartilaginous arc projects both dorsally along the caudal part of the tympanum and caudally in relation to the B2 pair. This pair is located in the dorsal part of the syrinx, in a transverse plane, each of which has round edges and a concavity towards the ventro-lateral part of the syrinx; a cartilaginous projection extends in the same shape into the caudal direction, almost reaching the B1 cartilaginous arc. Each of the B3 pairs is a cartilaginous arc-shaped half-ring whose concavity extends toward the lumen of the bronchus. Slightly medial in relation to the B1 arc, each pair has ventral and dorsal extremities that serve as attachments for one of the vibratory membranes.

### The syringeal muscles of the black jacobin

All syringeal muscles of the black jacobin are paired (Fig. 2). There is one pair of extrinsic, the *musculus tracheolateralis* (tracheolateral muscle; TL) and at least three pairs of intrinsic syringeal muscles: *musculus syringealis cranialis dorsalis ventralis* (ventro-dorsal cranial syringeal muscle; VDCrS), *musculus syringealis lateralis dorsalis ventralis* (ventro-dorsal lateral syringeal muscle; VDLS), and *musculus syringealis caudalis dorsalis ventralis* (ventro-dorsal caudal syringeal muscle; VDCaS).

**Fig 2.**
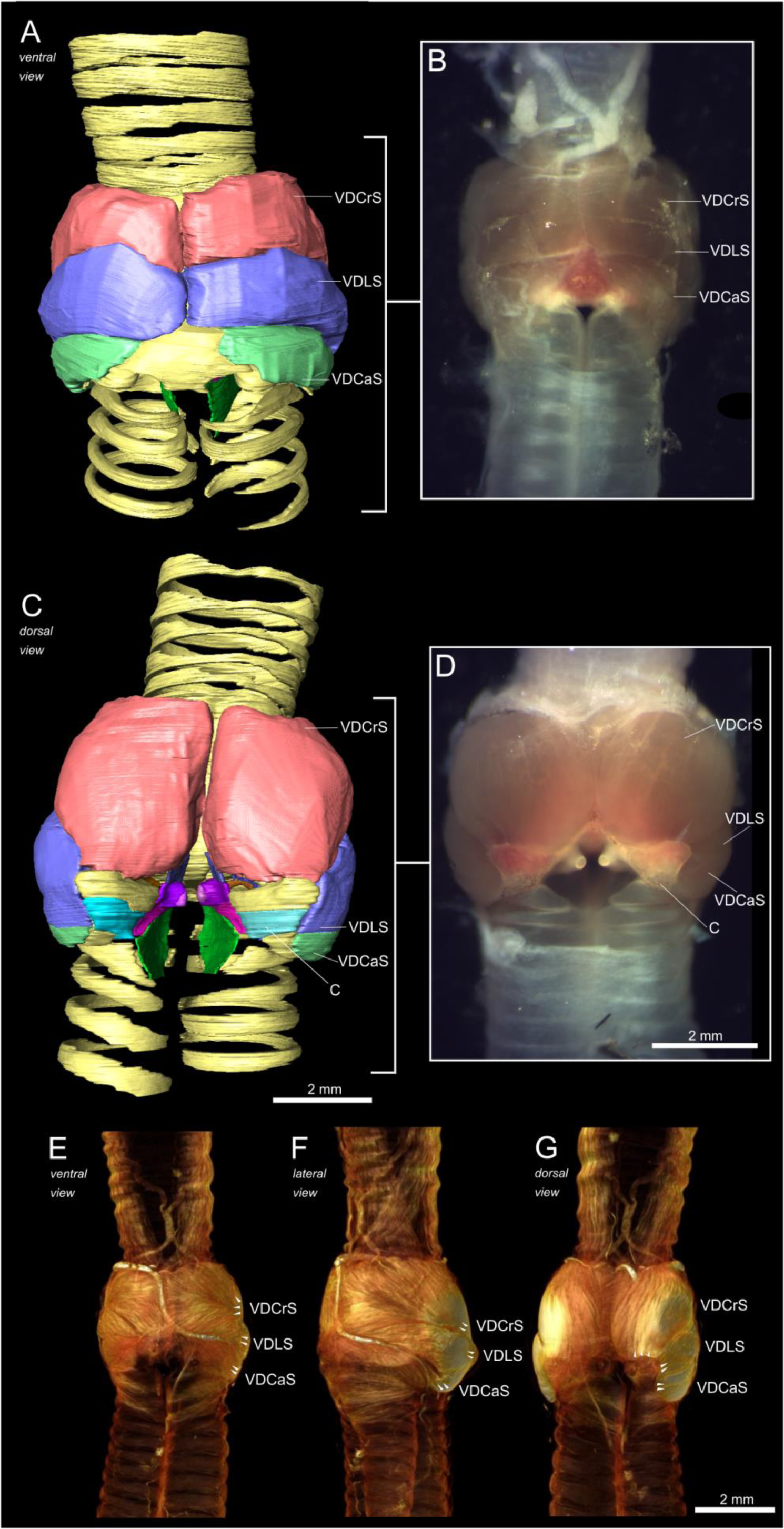
Musculature of the black jacobin’s syrinx. A and C, surface renderings; B and D, *in vitro* syrinx and E, F and G, volume renderings. A, B and E, ventral view; C, D and G, dorsal view and F, lateral view. The arrows indicate the orientation of the fibers. VDCrS, ventro-dorsal cranial syringeal muscle; VDLS, ventro-dorsal lateral syringeal muscle; VDCaS, ventro-dorsal caudal syringeal muscle and C, cartilage.

The extrinsic TL is composed of a few sparse sheets of muscle fibers attached to the cranial part of the trachea and extended caudally alongside the muscle’s lateral part until reaching the latero-cranial extremity of the tympanum (Fig. 2E–2G).

All intrinsic muscles are oriented ventro-dorsal. They attach ventrally to the tympanum and dorsally to some of the modified bronchial half rings. The VDCrS, the largest muscle, is responsible for 54% of the intrinsic musculature volume and follows the typical cardioid contour of the dorsal syringeal surface (Fig. 2C and 2D). The VDCrS caudal attachment site is in the ventrocranial head of the modified bronchial half-ring B2. A few muscle fibers of the VDCrS are attached via connective tissue to the tympanic ossicles. The VDLS comprises 34% of the intrinsic musculature volume placed mainly in the lateral part of the syrinx (Fig. 2F). The caudal attachment site of the VDLS is located on the lateral extent of half-ring B2 and includes its cartilaginous expansion. The VDCaS makes up the remaining 12% of the intrinsic musculature volume and runs mainly ventrally (Fig. 2A and 2B). The attachment sites of the VDCaS are located at the most caudal concavities of the pessulus and along the lateral outline of the cartilaginous extension of the half-ring B1.

### Vibratory tissue of the black jacobin’s syrinx

The syringeal vibratory tissues are composed of a pair of lateral labia (LL), each labium located in the lateral part of each side of the syrinx, and a pair of medial labia (ML) that continue into the medial tympaniform membrane (MTM). ML and MTM form the medial vibratory mass (MVM) in the medial part of the syrinx caudal to the tympanum (Fig. 3A). The LL is placed parallel to the ML and extends cranially over the tympanic lumen and caudally among the half-rings B1 to B3 (Fig. 3A). The LL has 45% of the volume of the ML and is ventrally attached to the pessulus, dorsally to B2 and laterally to the medial part of the B1 (Fig. 3A). The MVM constitutes a continuous mass of vibratory tissue reduced to 39% in thickness from the cranially located ML to the MTM. The ML is cranioventrally attached to the pessulus and dorsally attached to the half-ring B2. In the craniodorsal part, the ML thickens nearly 5-fold in an extension that embeds the tympanic ossicles and projects to the air passage (Fig. 3B and 3C). This extension, which comprises 37% of the total volume of ML and the tympanic ossicles comprise a further 6%, connects to a muscle via a thin ligament such as was reported previously for hummingbirds (Zusi, 2013). The MTM is ventro-dorsally attached to the medial edges of the bronchial half-ring B3.

**Fig 3.**
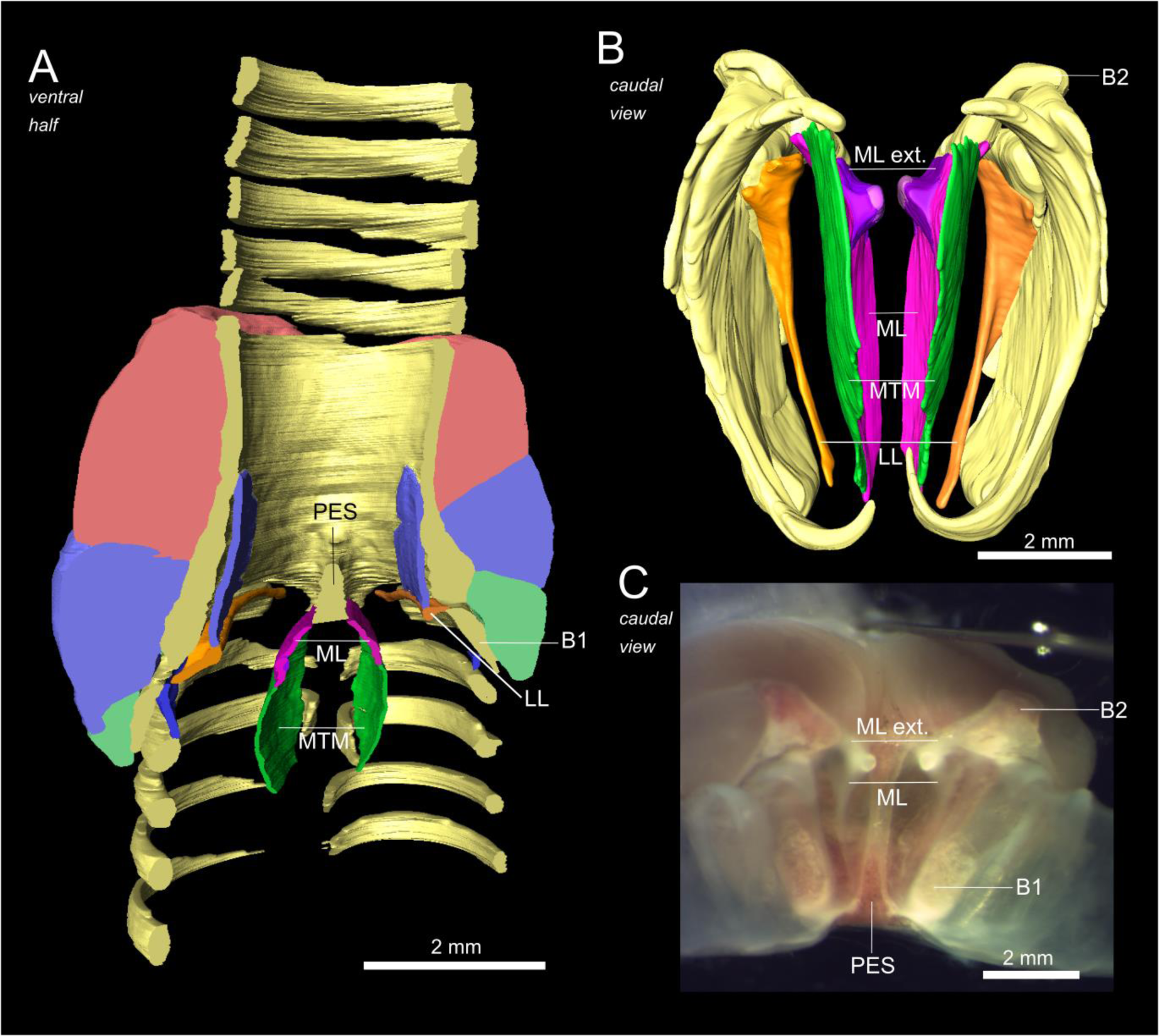
Vibratory membranes of the black jacobin’s syrinx. A and B, surface renderings and C, *in vitro* syrinx. A, ventral view and B and C, caudal view. In B, the first three pairs of bronchial half-rings and all pairs of vibratory membranes: LL (orange), MTM (green), ML (pink), ML extension (purple). PES, pessulus; ML, medial labium; MTM, medial tympaniform membrane; LL, lateral labium; B1, first bronchial half-ring; B2, second bronchial half-ring and ML ext., the cartilaginous extension of the ML.

### Biomechanics of the black jacobin’s syrinx

To explore potential general mechanisms of adduction, abduction, and stretching of the sound-producing elements, we carefully fixed one-half of the syrinx under a microscope and manually actuated the musculature around the ML *in vitro*. We identified the mechanism, which in sequential muscle activation, seems responsible for the adduction of the LL and stretching of the ML and LL. Applying an increasing amount of force to the caudal part of the cartilaginous arc extending from B1, the attachment site of VDCaS, resulted in an inward rotation of B1 and caused first partial and then complete adduction of the LL. A craniad force applied on the head of the B2, the attachment site of the VDCrS, resulted in the dorso-ventral stretching of the ML and LL (Fig. 4A). Considering the anatomical disposition of the VDLS, a lateral force applied on the lateral part of the B2, the attachment site of VDLS, may result in outward rotation of the B2 and cause the abduction of the ML. Thus, each of the three intrinsic muscles seems to be involved in one of the main tasks controlling sound production in the syrinx: the VDCaS on adduction, which closes the bronchial lumen; the VDLS on abduction, which opens the bronchial lumen; and the VDCrS, which controls the tension of ML and LL (Fig. 4B).

**Fig. 4.**
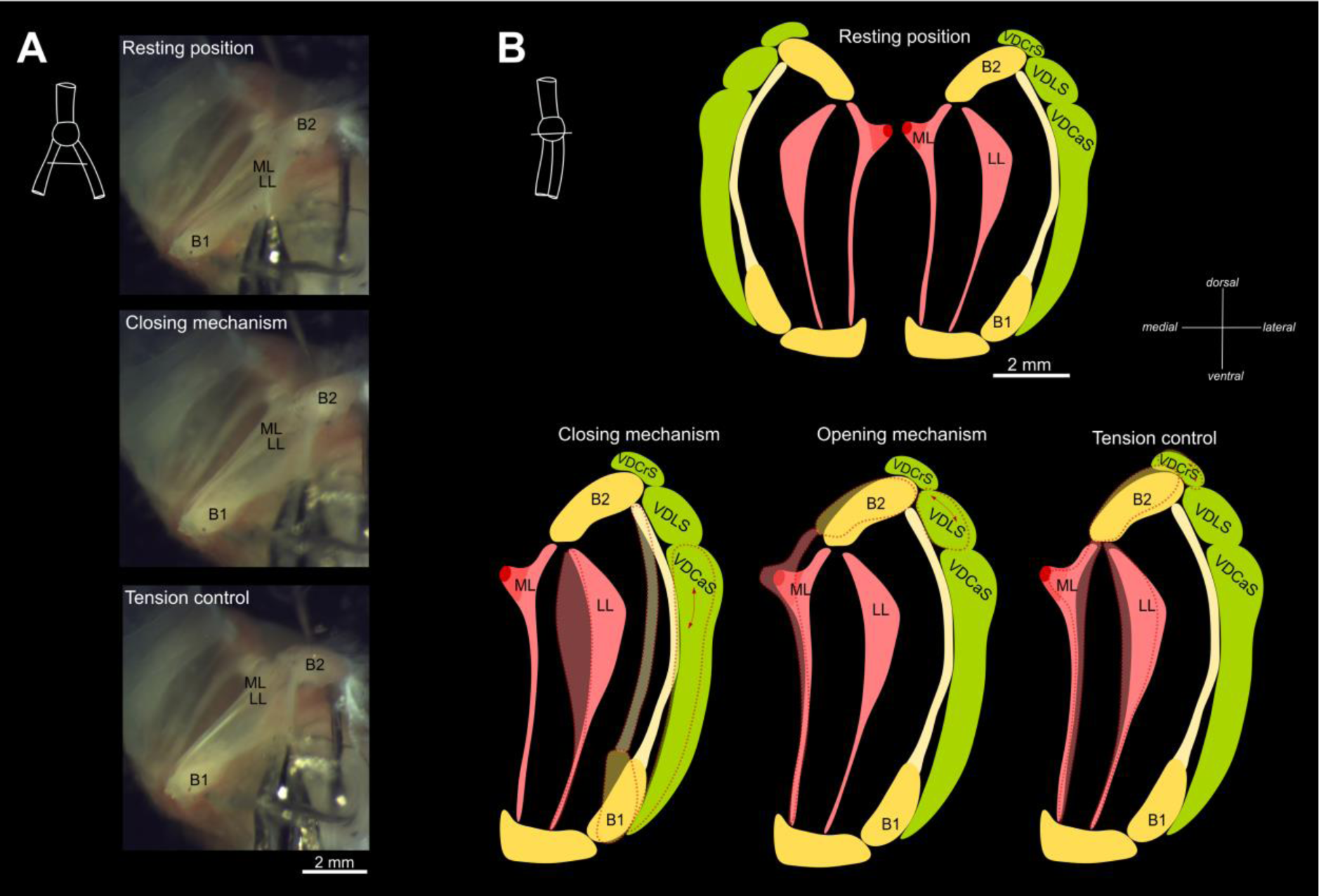
Biomechanics of the black jacobin’s syrinx. In A, caudal view *in vitro*. Upper panel: resting position; middle panel: inwards movement of the LL for abduction of the labia; lower panel: stiffening of the ML for tension control. In B, schematic representation of the syrinx, cranial view. Top: resting condition; bottom: closing, opening and tension control through hypothetical VDCaS, VDLS or VDCrS contraction. The shaded areas indicate the position of the syrinx in relation to the resting position and the red arrows indicate the involved muscle. ML, medial labium; MTM, medial tympaniform membrane; LL, lateral labium; B1, first bronchial half-ring; B2, second bronchial half-ring; ML, medial labia; LL lateral labia; VDCrS, ventro-dorsal cranial syringeal muscle; VDLS, ventro-dorsal lateral syringeal muscle; VDCaS, ventro-dorsal caudal syringeal muscle.

### Spectral analysis of the black jacobin’s vocalization

The black jacobin vocalizes with a fundamental frequency (F0) that ranges from 1.8 to 11.8 kHz (n = 105 recordings with a total of 1242 motor units, so-called syllables). We identified three types of vocalizations with distinct spectral structure: low-pitched vocalization with an F0 average of 1.8 kHz (± 0.5, n = 66 syllables); click-like chirps with an F0 average of 7.9 kHz (± 1, n = 148 syllables); and high-pitched vibratos with an F0 average of 11.8 kHz (± 0.4, n = 1028 syllables) (Fig. 5A).

**Fig. 5.**
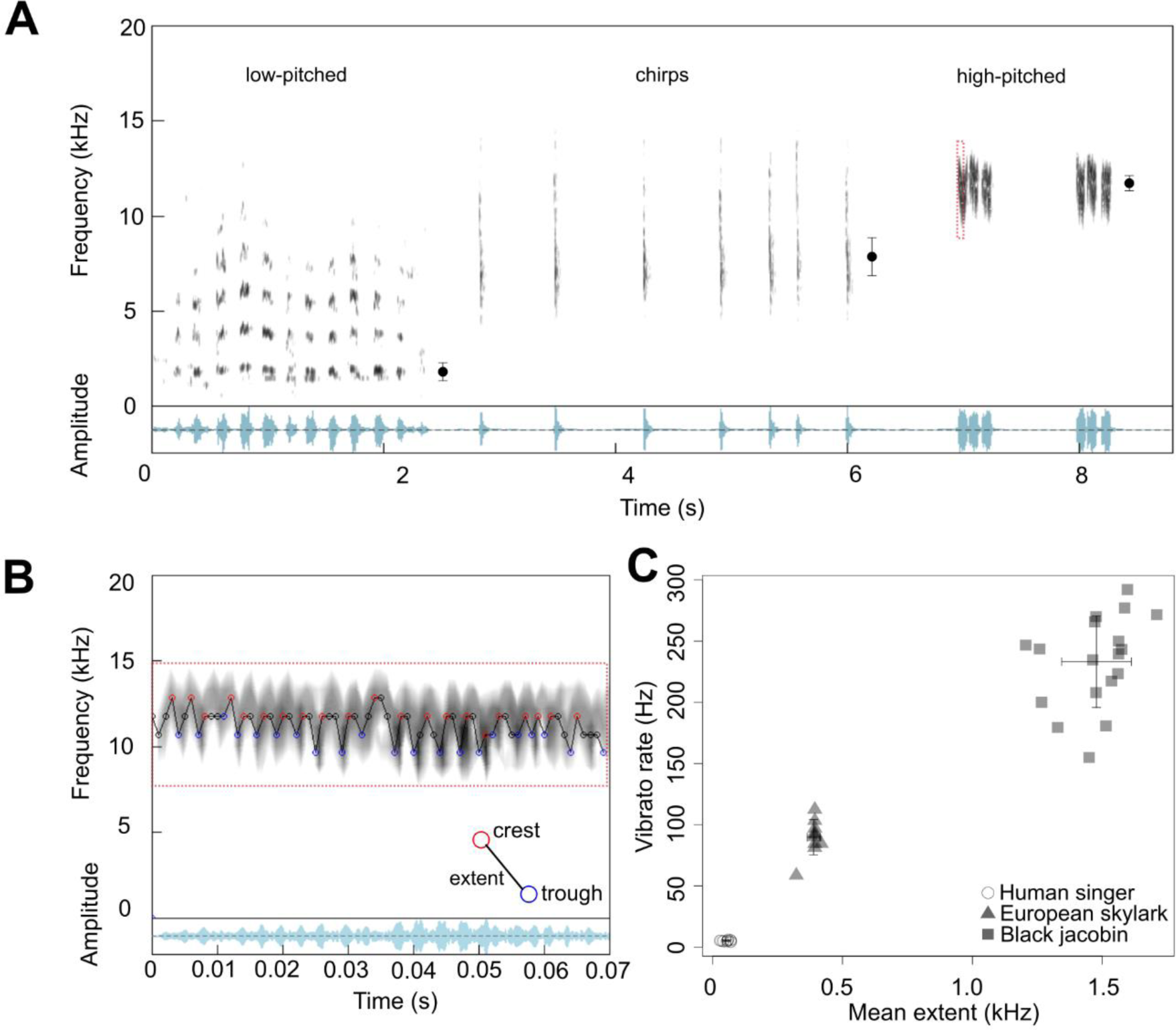
Black jacobin vocalizes in a wide frequency range with rapidly modulated vibratos. In A, spectrogram (frequency) and oscillogram (amplitude) of the three vocalization types produced by the black jacobin in the wild: low-pitched, chirps and high-pitched. The dots represent the average fundamental frequency for the respective vocalization on their left, and the whisker, the standard deviation. In B, an example of crest-trough detection in the high-pitched fragment is indicated by the box with red dashes in A obtained by our customized R function. Red circles indicate the crest, and blue the trough; the difference between them is the vibrato extent. In C, the vibrato rate is calculated as the number of crest and trough pairs per second given in Hz, and the mean extent for each syllable/segment is calculated in kHz. The cross indicates means and arrows the standard deviation. Triangles represent the data points obtained by an example of vibrato produced by a human singer (n = 12 vibrato segments in one song), the squares represent data points for the black jacobin (n = 18 syllables from three birds), circles for Eurasian skylark (*Alauda arvensis*) (n = 11 syllables from two birds). See the Methods for the source of the sound recordings.

The black jacobin’s most frequent type of vocalization is the high-pitched vibrato composed of syllables repeated in groups with up to three repetitions; the vibrato has a fundamental frequency of around 12 kHz with harmonics reaching the ultrasonic range of humans (this study; Olson et al., 2018). Each vibrato syllable is composed of fast oscillations with periodic changes of the fundamental frequency (Fig. 5B) and an average duration of 95.8 ± 35 ms (n = 18 syllables). Within a syllable, the difference between the highest and the lowest modulation of frequency, known as the vibrato extent, ranged from 0.7 to 3 kHz (average 1.5 ± 0.5 kHz, n = 401 crest-trough pairs, Fig. 5B) with a periodicity of around 2.4 ms (± 1.4 ms, n = 401). Thus, the black jacobin can produce syllables that change their fundamental frequency at an average rate of 233.2 Hz (± 37.5, n = 18 syllables). For example, Whitney Houston produced an average vibrato rate of 5.1 Hz (± 0.6, n = 12 vibratos) with a vibrato extent of 0.05 kHz (± 0.01) (Fig. 5C). This means black jacobin’s vibrato rate surpasses that of a human singer by more than 45-fold. Within birds, vibratos are reported to be produced by only a few species (Ferreira et al., 2006; Mundinger, 1995), but no quantification of vibrato rate or extent is available. For comparison, we quantified the vibrato rate of a songbird, Eurasian skylark (*Alauda arvensis*). The skylark’s vibrato rate is almost 17-fold higher than that of a human singer, with an average of 89.7 Hz (± 14.4, n = 10 syllables), and the vibrato extent averaged 8-fold larger, 0.4 kHz (± 0.02) (see Methods for details). No vibrato reported to date combines such a fast rate and wide extent as that of the black jacobin (Fig. 5C).

## Discussion

Here we present the first detailed description of the vocal tract of a basal hummingbird species, a species with the potential to illuminate how vocal learning has evolved. We identified the presence of heavily modified bones, intrinsic syringeal musculature with a particular orientation and a pair of vibratory tissues in each of the sides of the syrinx, each of which contains an ossified structure. This peculiar syringeal morphology allows the black jacobin to produce a vibrato that challenges the known limits of this acoustic feature.

### Syrinx stabilization and decoupling from physiological noise without ST muscles

The black jacobin’s syrinx is located outside of the thoracic cavity, in contrast to songbirds and parrots that have their syrinxes inside the thoracic cavity (Ames, 1971). Hummingbirds are highly specialized for hovering: unsurprisingly, its flight muscles make up 25 to 30% of its body weight, a ratio that is more than that of any other bird family (Schuchmann, 1999). The hummingbird’s enlarged flight muscles are combined with an enlarged heart, comprising about 2.5% of its body mass. The syrinx location outside of the thoracic cavity potentially alleviates spatial constraints and avoids mechanical disturbances from the enlarged flight or cardiac muscles. Since this syrinx displacement was reported in a few species of derived hummingbird sub-families (Beddard, 1898; Müller, 1878; Zusi, 2013), its occurrence in the black jacobin indicates a synapomorphy of the family. Such a syrinx location may have allowed the hummingbird to evolve control over its syringeal biomechanics by decoupling them from the harsh environment of internal organs.

In songbirds, the decoupling from internal organs is achieved through the sterno-tracheal muscle (ST), which is thought to stabilize the syrinx in most birds including the songbirds (Düring and Elemans, 2016; Suthers, 2004; Suthers and Zollinger, 2008). Black jacobins lack the ST altogether but seem to have further refined syrinx stabilization through tight wrapping of the syrinx in several layers of soft tissue. These layers create a rigid frame, protecting the syrinx from its immediate environment while keeping the syringeal elements inside flexible and maintaining the differential pressure necessary for the onset of sound production (Elemans et al., 2015; Gaunt and Gaunt, 1985). The most external of these layers might be an evagination of the interclavicular membrane that cranially encloses the syrinx within the interclavicular air sac, which has also been reported in other hummingbird species (Zusi, 2013).

In non-passerines, the ST is also involved in sound production. For example in the tracheal syrinx of pigeons, the shortening of the ST brings its cartilages closer together, thereby closing the syringeal lumen (Mindlin and Laje, 2006). The adduction of the labia is crucial for sound production in general as it allows to build up the phonation threshold pressure (PTP), which is necessary for sound onset (Titze, 1992). In songbirds, adduction is achieved by intrinsic musculature, rather than ST adduction (Elemans et al., 2015) Similarly, due to the lack of ST muscles, the closing mechanism in black jacobins is operated by intrinsic musculature.

The syrinx displacement may also have had implications for muscle orientation. The intrinsic muscles of the black jacobin’s syrinx are oriented dorso-ventrally, likely a synapomorphy of hummingbirds, while the intrinsic muscule fibers of most bird taxa run cranio-caudally (King, 1989; Smolker, 1947). Because all of the black jacobin’s intrinsic muscles are ventrally attached to the tympanum, but each of them is dorsally attached to a different point, they run dorso-ventrally on different angles. The general dorso-ventral orientation with differences in angulation might allow the black jacobin to control the mobile syringeal elements despite the lack of lateral stabilization provided by the STs in other taxa.

The gradual disposition of the syrinx and accompanied general absence of STs in hummingbirds (Müller, 1878), might have been one of the driving pressures for the evolution of intrinsic muscles, a key prerequisite of vocal learning.

### Tympanic ossicles

Although cartilaginous formations were found embedded in the vibrating tissues of songbirds (Düring et al., 2013), tympanic ossicles have not been reported in any species other than hummingbirds (Müller, 1878; Zusi, 2013). The origin of tympanic ossicles is uncertain. Due to their medial position and proximity to the tympanum, they might be either modified bronchial half-rings or have originated from a tracheal ring. In humans, the prevalence of a small sesamoid bone in the knee has increased worldwide in the past century, probably as a dissipative response to increased mechanical forces due to the enlargement of leg bones and muscles (Berthaume et al., 2019). Similarly, increased tension in the labia might have led to the formation of tympanic ossicles in the black jacobin’s syrinx.

In addition to direct muscular activity, stiffness of vocal tissues depends on the elastic properties of the tissue itself (Düring et al., 2017; Riede et al., 2010). In songbirds, cartilage embedded in the medial labia (ML) both aids in the dissipation of the tension, avoiding rupture under high stress and modifies the elastic properties of the syrinx (Düring et al., 2013). In particular, the cartilage that connects with the muscle potentially supports a more gradual bending mechanism, which in turn allows uncoupling the control of amplitude and frequency (Düring et al., 2013). Similarly, this might be the function of the cartilaginous extension in the dorsal part of the ML and its embedded ossicles, the tympanic ossicles, in the black jacobin. This extension is connected by a thin strip of connective tissue to a few muscle fibers of the larger syringeal muscle; given this arrangement, direct muscular control of the extension seems likely.

The tympanic ossicles may contribute to achieving the black jacobin’s high fundamental frequency: they cause high local density and prevent an entire part of the ML from vibrating at all, thus shortening its length and increasing the fundamental frequency. In other words, the tympanic ossicles could be used as a secondary mechanism to gradually increase ML stiffness and reduce ML length. It is therefore likely that the cartilaginous extension of the ML in the black jacobin both shifted the elastic properties of the ML towards the optimal for high fundamental frequency by increasing ML density towards the muscle attachment site that directly controls ML stiffness, and shortened the vibratory part of the ML.

### Extreme vocal performance

Black jacobins produce particularly rapidly-modulated vibrato sounds (this study; Olson et al., 2018). The black jacobins’ vibratos oscillate periodically up and down with a frequency bandwidth of up to 3 kHz at a rate of about 250 Hz. This fast vibrato rate can be compared to that of other extreme vocal performances, such as of starlings (*Sturnus vulgaris*), a songbird whose muscle activity in the syrinx produces changes in sound amplitude at a repetition rate of 218 Hz (Elemans et al., 2008). The musculature of the songbird’s syrinx belongs to a special class of muscles, called superfast muscles (Elemans et al., 2004; Rome, 2006), and can produce work at cycling limits of approximately 90 Hz to 250 Hz (Rome et al., 1996). *In vitro* preparations revealed that the superfast songbird muscles in the syrinx have the potential to function at cycle frequencies as fast as 250 Hz (Elemans et al., 2008). Although direct electromyographic recordings of the syringeal musculature would be needed to confirm that the black jacobin’s vibrato rate of 250 Hz is a direct result of muscular control, this extremely fast performance suggests that the black jacobin’s syringeal muscles produce work on the upper limit of the superfast muscle activity reported to date (Elemans et al., 2008) and that black jacobins may have muscle properties comparable to those of songbirds.

### Biomechanics of sound production and implications for vocal learning in hummingbirds

Parrots and songbirds, two vocal learners, have a tracheal and a tracheobronchial syrinx, respectively, both with intrinsic musculature (Ames, 1971; Düring et al., 2013; Gaban-Lima and Höfling, 2006; Nottebohm, 1976). The black jacobin’s syrinx, like that of all the other hummingbird species reported so far (Müller, 1878; Zusi, 2013), is tracheobronchial, with three pairs of intrinsic muscles that are as complex as those of songbirds. The black jacobin’s multiple intrinsic muscles attach in close proximity to movable elements of its syringeal skeleton (modified bronchial bones) to which the vibrating tissues (medial labia or lateral labia) are attached via cartilaginous extensions. These muscles seem to operate constitutively. For example, both lateral and medial labia are attached to the bronchial half-ring B2, where two large muscles are attached. At its cranial surface is the ventro-dorsal cranial syringeal muscle (VDCrS), and at its lateral part, the ventro-dorsal lateral syringeal muscle (VDLS). Various amounts of contraction of each muscle might contribute gradually to distinct functions, such as the abduction of the ML and the stretching of the labia. Since position and tension of the labia are directly related to distinct acoustic parameters, multiple muscles contributing to the same function creates redundancy in possible motor commands controlling accoustic parameters such as pitch. When the brain has multiple, rather than a unique, motor command available to achieve a certain vocal output, a redundant control space may simplify trial-and-error attempts during imitation in the vocal production learning process (Elemans et al., 2015).

## Conclusion

Here we present the first high-resolution morphology of a hummingbird syrinx, the black jacobin’s. We suggest the absence of sternotracheal muscle, presence of tympanic ossicles, dorso-ventral muscle orientation and syringeal displacement as synapomorphies within hummingbirds. These characteristics might have evolved concomitantly with the displacement of the syrinx out of the thorax, as an operational solution to reduce interference of the syrinx with the enlarged heart and flight muscles. The biophysical and biomechanical redundancies emerging from the hummingbird’s syrinx morphology may represent a crucial step towards the evolution of vocal learning in hummingbirds.

## Methods

### Tissue collection and preparation

The black jacobins (n = 3) were captured with a hummingbird-specific “Ruschi trap” (Ruschi, 2009) in the park of the National Institute of the Atlantic Forest (former Professor Mello Leitão Museum), Espírito Santo State, Brazil, in accordance with the Brazilian Institute of Environment and Renewable Natural (IBAMA) regulations under the Biodiversity Information and Authorization System (SISBIO) license number 41794-2.

Two males were deeply anesthetized with ketamin (15mg/kg) and perfused through a cardiac injection with the following sequence of solutions: 0.5 ml heparin-nautriun anticoagulant, 0.9 saline buffer and 4% paraformaldehyde fixative. After the perfusion, the syrinx was dissected and stored in the fixative for 24 hours and then stored in 0.1 M phosphate-buffered saline (PBS) in solution with 0.05% sodium acid until use. We used both fixed syrinxes for micro-computed tomography, one stained for the visualization of soft tissues and the other without the staining procedure for clear visibility of the ossified structures. Both syrinxes were dissected with a large part of the esophagus and bronchi as close as possible to the beak and lungs, respectively, to access the syrinx structures integrally. A third male black jacobin was killed with an anesthetic overdose and the syrinx immediately dissected and cryopreserved at - 80 °C until use. This syrinx was micro-dissected.

### Micro-dissection

The cryopreserved syrinx was thawed gradually. First, at −20 °C for one hour followed by 24 °C during the time of manipulation. For manipulation, the syrinx was pinned down on a glass Petri dish covered by black dissecting pan wax and filled with 0.1 M PBS. We disassembled the syrinx under an MZ75 stereomicroscope (Leica Microsystems, Germany) equipped with an ISH500 5.0 MP camera (Tucsen Photonics, China).

The syrinx was inspected ventrally and dorsally; the main musculature and ossifications matched with the μCT-based reconstitution. The difference in the density of adjacent soft tissues was noted by a comparison of their light reflection. We sectioned the muscles at their tympanic insertion site and noted the general orientation of fibers. We repositioned the syrinx caudally, centering it where the bronchia bifurcated, with bronchi angled at 180° exposing the vibratory tissues. The mobile structures in which the vibratory tissue was attached were noted. With a pin, we applied gentle force to each of these mobile structures and photographed the effect of the applied force on the vibratory tissue.

### Micro-computed tomography

The micro-computed tomography (μCT) scans of isolated syrinxes (two males) was conducted at the Zoologische Staatssammlung München (Munich, Germany) using a phoenix nanotom m cone beam µCT scanner (GE Measurement and Control, Wunstorf, Germany) with down to 3.1 µm voxel size.

One syrinx was scanned without staining to access the anatomy of the ossified structures as a fourfold multiscan with the following parameters: 100 kV source voltage, 170 μA source current, 0.1 mm aluminum filter, 500 ms exposure time, 3.1 μm isotropic voxel resolution, 1000 projections over 360° with three averaged images per rotation position, and a total of 132 min scan time, using a molybdenum target. The second syrinx was stained with a contrast agent to image soft tissues. It was placed inside a glass vial with 0.1% Lugol’s solution (Sigma Aldrich). The vial was placed on a tube roller for 48 h. The stained syrinx was scanned for 48 minutes using the following parameters: 80 kV source voltage, 180 μA source current, 0.1 mm copper filter, 500 ms exposure time, 3.6 μm isotropic voxel resolution, 1440 projections over 360° with three averaged images per rotation position, using a tungsten (“standard”) target. The volume reconstructions were performed using the software phoenix datos׀x 2 (GE Sensoring & Inspection Technologies GmbH, Germany).

### Three-dimensional reconstruction and nomenclature

The annotation was performed onto the μCT-based syringeal dataset of a black jacobin adult male. We identified the recognized musculature, ossification, cartilaginous pads, and vibratory tissues. The visualization procedures including volume rendering and manual segmentation for surface rendering, and relative quantifications were done with the software Amira 6.1 (Thermo Fisher Scientific, Massachusetts, USA).

The syrinx structures were defined by the consensus of the microdissection and the µCT data. The nomenclature was given following the same procedure used in Düring et al. (2013) (Table 1). We named the tracheal rings T1 to Tn starting from tympanum and moving toward the larynx. We present a conservative number of intrinsic muscles due to their delineation. We delineated the intrinsic syringeal muscles by aggregating fibers that were oriented at the same angle and defined their differential attachment sites based on both microdissection and the μCT scans. The extrinsic musculature was not traced in the 3D reconstruction due to its undetermined tracheal insertion but is partially shown. The vibratory tissues we found are analogous to those described in Düring et al. (2013).

**Table 1.**
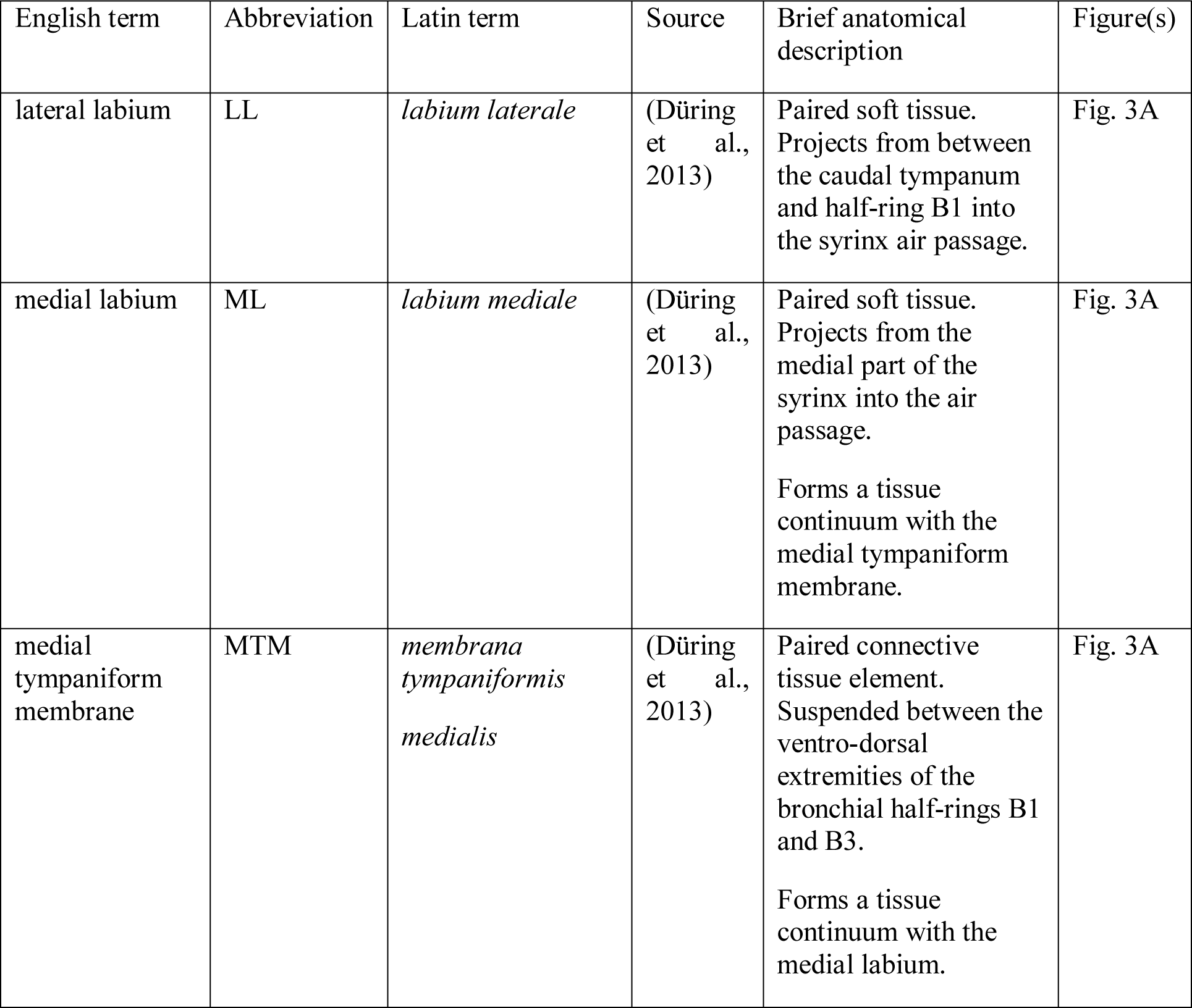
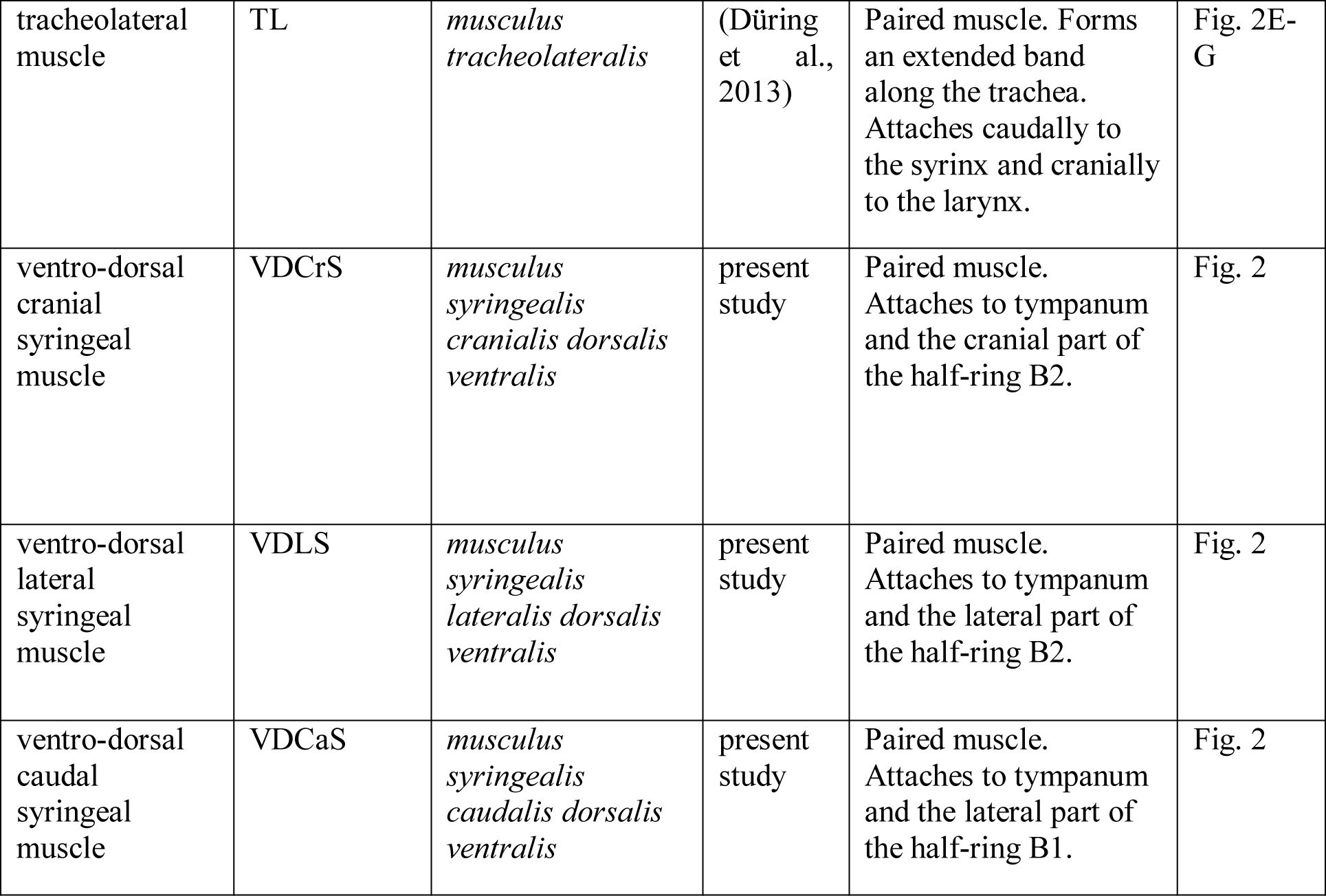
Anatomical structures of the black jacobin syrinxes described in this study. The source indicates when the name is given in analogy with a previous description for a different species

### Sound analysis

First, we investigated the fundamental frequency of the black jacobin’s vocalizations in the wild. Observations and recordings were made in the Professor Mello Leitão Museum (Santa Teresa, Espirito Santo state, Brazil) at a feeding point where every day wild hummingbirds visit feeders that contain 25% sugar water solution. The observations were made over nine days from November to December 2013, and over 15 days from September to October 2015 during the black jacobin’s breeding season (Ruschi, 1964). Black jacobins were observed continuously for one hour a day on the dates mentioned above; observations were made sometime between 6:30 and 11 a.m. for a total of approximately 24 hours. The sampling method was *ad libitum* (Martin and Bateson, 2007), according to which the most conspicuous occurrences of the vocal behavior were recorded for the first black jacobin spotted at the feeding point until the individual had left the place. The black jacobins were not individually marked, but the high abundance of the species at the feeding point (Loss and Silva, 2005) and the fact that recordings were obtained over two non-consecutive years make it unlikely that the observations were biased toward a few individuals. Recordings were made 3–10 m from the individuals with a *Marantz PMD 671* (Marantz, New York, USA) solid-state recorder connected to a Sennheiser MKH 70/P48 (Sennheiser, Wedemark, Germany) directional microphone in a 48 kHz sampling rate wave file. We obtained 105 recordings totaling five hours. We isolated the black jacobin’s vocalizations and calculated the fundamental frequency for each of their syllables (vocal units) using the packages “Seewave” (Sueur et al., 2008) and “WarbleR” (Araya-Salas and Smith-Vidaurre, 2017) in R 3.5.0 (R Core Team, 2018). The recordings are not public due to storage reasons but are available from the corresponding author upon request.

Second, we focused on the most common vocalization of the black jacobin. This vocalization is composed of syllables with continuous and regular fast modulations in fundamental frequency (Olson et al., 2018). Given the periodicity of these modulations, we classified the syllables as vibratos. Vibrato is a demanding vocal task produced by opera singers and characterized by periodic pitch fluctuation (Sundberg, 1994). The accuracy of the vocal performance can be quantified in terms of four parameters: rate, extent, regularity, and waveform (Sundberg, 1994). Here we measured two features of the black jacobin’s vibrato: the rate that was measured by the number of oscillations per second and the extent that was the depth of the oscillations. We measured the vibrato based on and adapted from Migita et al. (2010). All calculations were performed on the platform R 3.5.0 (R Core Team, 2018). For the calculations, syllables were selected from full recordings using the package “WarbleR” (Araya-Salas and Smith-Vidaurre, 2017), then the fundamental frequency contour of each unit was identified with “Seewave” (Sueur et al., 2008), and crest-trough pairs were detected using a customized R script. The vibrato rate given in Hz was calculated by:

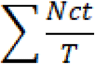

Where Nct is the total number of crest-trough pairs detected per vocal unit, and T is the total duration of the unit in seconds.

The vibrato extent given in Hz was obtained per crest-trough pair by the difference between the frequency of the crest and the frequency of the trough. The values were presented as means (± standard deviation, sample size).

We analyzed three high-quality sound recordings and 18 syllables with the highest quality obtained from three black jacobins. These recordings were kindly provided by Olson et al. (2018), who obtained them using an Avisoft CM16/CMPA ultrasound microphone (2– 250 kHz range) coupled to an UltraSoundGate 416H amplifier recorder at the frequency rate of 214 kHz. To have something to compare with the black jacobin, we analyzed the soundtrack “I will always love you” performed by Whitney Houston (© Sony Music, 1992) and selected 12 fragments in which the singer produces a vibrato as an example of a human singer. As an example of a vibrato produced by a songbird, we analyzed two recordings of the Eurasian skylark (*Alauda arvensis*) obtained from the Xeno-canto collaborative bird sound collection (https://www.xeno-canto.org/), catalog numbers XC401962 and XC417772 uploaded by Karl-Birger Strann and Jarek Matusiak, respectively. The vibrato examples of both the human singer and songbird were analyzed following the same parameters as the black jacobin recordings.

## Acknowledgments

We are grateful to the staff of the Museu de Biologia Professor “Mello Leitao” (current National Institute of the Atlantic Forest) for assistance with data collection in the field and the former director Hélio de Queiroz Boudet Fernandes for logistical support. We thank Christopher Olson for providing high-quality sound recordings of the black jacobin. AM received an IMPRS Organismal Biology grant for language editing; we thank E. Wheeler for the helpful comments.

## Authors’ contributions

AM and DD conceived the study. AM, AC, BR, and DD contributed to data acquisition. DD, AC, BH, and MG contributed reagents, materials, and analytical tools. All authors contributed to data analysis. AM wrote the initial draft of the manuscript. All authors contributed to manuscript revision and approved the final version.

